# Population coding of distinct categories of behavior in the frontal eye field

**DOI:** 10.1101/2023.11.28.568997

**Authors:** Matan Cain, Mati Joshua

**Affiliations:** Edmond and Lily Safra Center for Brain Sciences, The Hebrew University of Jerusalem, Jerusalem, Israel

## Abstract

Brain regions frequently contribute to the control of diverse behaviors. Within these regions, individual neurons often respond to various motor tasks, suggesting that the ability to control multiple behaviors is rooted in the organization of neuronal populations. To study this organization, we examined how the frontal eye field (FEF) encodes distinct behaviors by recording the activity of 1200 neurons during various eye movement tasks in two female fascicularis monkeys. We focused on three behaviors: smooth pursuit, pursuit suppression, and saccades. We found that single neurons tended to respond in all three tasks, thus challenging the notion that the FEF is organized into task-specific clusters. We then identified the low-dimensional subspaces that contained most of the population activity during each behavior and quantified the extent of overlap between these spaces across behaviors. Population activity during pursuit and pursuit suppression exhibited a substantial overlap, with highly correlated directional tuning at the single-neuron level, as reflected by similar outcomes for the population decoders. These distinct behaviors combined with similar encoding suggest that the suppression of movement occurs mostly downstream from the FEF. By contrast, pursuit and saccades mostly occupied orthogonal subspaces, prompting an independent linear readout of saccades and pursuit. Thus overall, these results indicate that distinct behaviors can exhibit either separate or overlapping population codes within a specific brain region, hence emphasizing the importance of the system-level organization of behavior.

**Significance statement:** How do brain areas control multiple behaviors? We investigated the organization of the FEF in monkeys during three different types of eye movements: smooth pursuit, pursuit suppression, and saccades. We found that individual neurons tended to respond to all three tasks, thus challenging a task-specific FEF cluster organization. Further, the low-dimensional subspaces that contained most of the population activity during pursuit and pursuit suppression overlapped substantially, implying that movement suppression occurs downstream from the FEF. In contrast, pursuit and saccades occupied orthogonal subspaces, prompting independent linear readouts. These results underscore the importance of adopting a system-level perspective to comprehend how diverse behaviors are encoded in the brain.

## Introduction

The brain processes a multitude of stimuli to control a wide range of behaviors. To achieve this goal, brain regions, rather than exhibiting strong specificity by solely encoding one stimulus or controlling a singular behavior, frequently process multiple stimuli and generate a diverse set of behaviors. This lack of specificity underscores the need to examine the underlying organization at the population level responsible for encoding specific sensory inputs and generating particular behaviors. Consider for instance the challenge of generating distinct categories of movements within a singular brain region. Do the same neurons encode distinct behaviors? If so, how is the network organized within and across populations to produce different patterns of movement?

An experimentally trackable and well-studied example of the control of multiple behaviors can be found in the FEF, a central node in the eye movement system (Bruce and Goldberg, 1985; Lynch, 1987; Macavoy et al., 1991; Schall, 1991, 2004). The FEF controls categorically distinct behaviors such as saccades (Bruce and Goldberg, 1985; Schall, 1991) which are characterized by brisk, rapid eye movements, as well as smooth pursuit (Lynch, 1987; Macavoy et al., 1991; Gottlieb et al., 1993, 1994), the prolonged, slow eye movements driven by visual motion. The FEF also plays an important role in inhibiting both saccades (Hanes et al., 1998; Brown et al., 2008) and smooth pursuit (Fukushima et al., 2011). Numerous studies have examined the relationship between the activity of individual neurons in the FEF and behavior (Bruce and Goldberg, 1985; Tanaka and Lisberger, 2001; Schall, 2004; Costello et al., 2013; Raghavan and Joshua, 2017). These studies often focus on activity during either saccades or pursuit. However, a substantial part of all FEF neurons are involved in both eye movements (Mayo, 2022). The absence of task-specific clusters, along with the multitasking properties of FEF neurons raises the question of how saccade and pursuit are encoded at the population level.

Recent methodological advances now provide new techniques for dissecting the population organization that cannot be directly inferred from the activity of individual neurons (Churchland et al., 2012; Kaufman et al., 2014; Elsayed et al., 2016; Gallego et al., 2018). These techniques characterize the subspace of activity that is spanned by the population activity and the trajectory of the activity in these spaces. The relationship between the subspaces defined by population activity during different eye movements has significant implications for reading out the information from the population activity. For example, an orthogonal organization would enable an independent readout of specific categories of eye movements, whereas a substantial overlap would require further mechanisms to distinguish between behaviors (Kaufman et al., 2014; Elsayed et al., 2016).

In this study, we used population analysis methods to explore the organization of the FEF in three distinct eye movement tasks: smooth pursuit, suppression of pursuit and saccades. We used multi-contact probes to record a large number of neurons simultaneously, thereby reducing the sampling bias that can arise from selecting neurons based on their response properties. The findings reveal that the neural activity patterns during smooth pursuit and suppression tasks were largely similar suggesting that the distinction between pursuit and suppression occurs downstream from the FEF in the eye movement neural pathway. Conversely, the subspace defined by the activity in the saccade task was mostly orthogonal to the subspaces spanned in the pursuit and suppression tasks. Collectively, the findings show that distinct behaviors can be encoded in a specific brain region in either a similar or dissimilar manner, hence highlighting the importance of the system-level organization of movement.

## Methods

We collected neural and behavioral data from two females (monkey F and monkey J) Macaca Fascicularis monkeys (4-5 kg). All procedures were approved in advance by the Institutional Animal Care and Use Committees of the Hebrew University of Jerusalem and were in strict compliance with the National Institutes of Health Guide for the Care and Use of Laboratory Animals. We first implanted head holders to restrain the monkeys’ heads in the experiments. After the monkeys recovered from surgery, they were trained to sit calmly in a primate chair (Crist Instruments) and consume liquid food rewards (baby food mixed with water and infant formula) from a tube set in front of them. We trained the monkeys to track spots of light that moved across a video monitor placed in front of them.

Visual stimuli were displayed on a monitor (60 cm from the eyes of the monkeys). The stimuli appeared on a dark background in a dimly lit room. A computer performed all real-time operations and controlled the sequences of target motions. The position of the eye was measured with a high temporal resolution camera (1 kHz, Eyelink 1000 plus, SR research) and collected for further analysis. We performed a second surgery to place a round 19 mm diameter recording cylinder over the FEF. The center of the cylinder was placed above the skull at 19 mm anterior and 15 mm lateral to the stereotaxic zero, on the left hemisphere for monkey F and on the right for monkey J. Localization of the FEF was based on stereotactic coordinates, the post-implant MRI scan and initial mapping with a single contact electrode. In addition, we confirmed the FEF location with electrical stimulation (Bruce et al., 1985). Before (Monkey J) or after (Monkey F) the recording sessions, we evoked eye movements by applying electrical currents (24 biphasic pulses, 50 μA, 350 Hz).

After identifying the FEF in each recording session, we inserted either a 24-channel (monkey F) or a 64-channel (monkey J) linear probe (Plexon S-probes with a contact spacing of 150 or 125 µm) into the FEF. When lowering the probe, we aimed to maximize the number of neurons recorded.

Signals were digitized at a sampling rate of 40 kHz (OmniPlex, Plexon). For the detailed data analysis, we sorted spikes offline (Plexon). For sorting, we used principal component analysis and corrected manually for errors. We visually inspected the waveforms in the principal component space and only included neurons for further analysis when they formed distinct clusters from noise. However, some clusters did not have clear boundaries in the PC space (estimated visually as 664/1213). We therefore confirmed that including only well-isolated clusters did not alter the results. Specifically, the overlap in tuning between tasks persisted for well-isolated neurons. Crucially, we applied the same sorting criteria consistently across all tasks, so that any differences between tasks cannot be attributed to the sorting process. Finally, we primarily used population measures that are insensitive to sorting errors (Trautmann et al., 2019). We converted the sorted spikes into time-stamps with a time resolution of 1 ms and conducted a visual inspection once again to check for instability and rectify obvious sorting errors.

### Experimental design

Pursuit task: Each trial started with a bright white circular target that appeared in the center of the screen. After 500 ms of presentation, in which the monkey was required to acquire fixation, a target appeared at a position 4° eccentric to the fixation target at one out of eight equally spaced directions. We defined this time point as the cue onset. At this point, monkeys were required to maintain fixation on the central target, and moving the eye towards the eccentric target would result in failure. After a variable delay of 800-1200 ms, the fixation target disappeared and the eccentric target moved towards and through the center of the screen at 20°/s. The target moved for 750 ms, then stopped and stayed still for an additional 300-500 ms. If the monkey’s gaze was within a 3-5°x3-5° window around the target, the monkey received a reward.

Suppression task: The structure of the suppression task was identical to the pursuit task, except that the target at the center of the screen was a square and it did not disappear at motion onset. After motion onset, the monkey was required to maintain fixation within a 3-5°x3-5° window around the central target rather than on the eccentric moving target.

Saccade task: Each trial started with a bright white circular target that appeared in the center of the screen. The monkey had 500 ms to acquire fixation and then had to maintain fixation for a variable delay of 800-1200 ms. The central target disappeared and immediately reappeared in one of eight equally spaced eccentric locations 10° from the center of the screen. If the monkey’s gaze was within a 5°x5° window around the target, the monkey received a reward.

### Data analysis

Analyses were performed using MATLAB (MathWorks) or Python

To quantify the directional tuning of a neuron over time, we calculated the partial *ω*^2^ effect size (*ω*^2^_p_). *ω*^2^_p_ is a common effect size measure used in ANOVA designs that is often preferred over other effect size measures since it is unbiased (Olejnik and Algina, 2000, 2003; Okada, 2013; Larry et al., 2022). We counted the number of spikes in bins of 100 ms, resulting in a distribution of the number of spikes emitted by the neuron in each condition and each bin. The calculation is as follows:

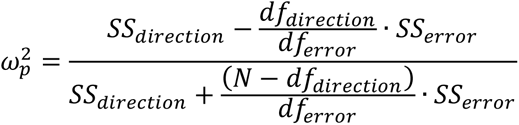

where *SS_direction_* is the ANOVA sum of squares for the effect of the direction, *SS_error_* is the sum of squares of the errors after accounting for all experimental variables, *df_direction_* and *df_error_* are the degrees of freedom for the direction and the error respectively and *N* is the number of observations (number of trials).

To calculate the coding of directional tuning throughout the entire epoch, we calculated *ω*^2^ in a model that included time as an additional variable. Again, we calculated the number of spikes in 100 ms bins, from 0 to 800 ms after an event (cue or motion onset). We fitted an ANOVA model that included the direction of the trial, with the addition of time (the specific bin the sample came from), and included interactions between time and direction. The formula we used was:

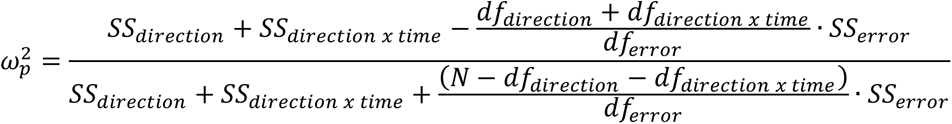

where *SS_direction x time_* is the ANOVA type II sum of squares for the interaction, and *df_direction x time_* is the corresponding degrees of freedom. We included the interaction term to quantify the time-varying coding of the direction. In this case, *N* is equal to the number of trials times the number of time bins. We excluded the main effect of time since we were interested in quantification of the directional tuning.

The *ω*^2^_p_ takes into account modulations across conditions and the trial-by-trial variability of the neuron. This contrasts with other effect size estimators based solely on the average activity, which can be strongly biased by the trial-by-trial variability. The *ω*^2^_p_ also has the advantage that it can be used with multiple conditions (such as different directions of movement) unlike other measures such as the choice probability that are defined for two task conditions (Larry et al., 2022).

We determined the preferred direction of each neuron as the closest direction to the direction of the vector average of responses within the first 350 ms after the relevant event across all directions, essentially representing the direction of the center of mass. Using shorter time windows (250 ms) did not alter any of our conclusions.

To examine the average time-varying properties of the response, we calculated the peri-stimulus time histogram (PSTH) at a 1 ms resolution. We then smoothed the PSTH with a 20 ms standard deviation Gaussian window, removing at least 100 ms from the edges to avoid edge effects. For analyses involving covariance, correlation or PCA, we mean-centered the neural activity for each neuron in each task. This was achieved by subtracting the mean activity across all conditions for each neuron at each time point. We then concatenated these PSTHs. For these analyses we used the 350 ms after the event. Taking shorter time intervals (250 ms) did not alter any of our conclusions.

To calculate the tuning consistency score for each pair of neurons, we computed the covariance of the concatenated PSTHs. We chose to use covariance rather than the correlation coefficient because it is closely related to principal components analysis. Note that using the correlation coefficient did not affect any of our conclusions. To assess whether the correlation in tuning covariance between the pursuit and suppression tasks (r_1_) was significantly different from the correlation between the pursuit and saccade tasks (r_2_), we first determined the z values (z_1_ and z_2_) corresponding to the correlation coefficients using Fisher’s r to z transformation (Fisher, 1915) and defined Z_observed_ as:

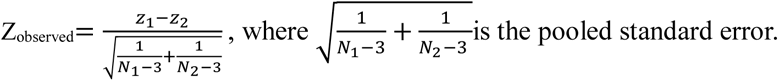

Finally, we assessed statistical significance by determining whether Z_observed_ was greater than the critical value (0.05).

To perform principal component analysis (PCA), we constructed for each task a population activity matrix A_cond_ ∈ R^N*CT^ where N represents the total number of neurons, C the number of directions and T the number of time points in each PSTH (350). To identify the principal components, we designed a new matrix P_cond_∈ R^N*CT^ based on the population activity matrix, where the activity at each time point was mean-centered by subtracting the average value from each column. The principal components for a given direction, PCs_cond_ ∈ R^N^, were defined as the eigenvectors of the covariance matrix of P_cond_. Using shorter time windows did not alter any of our conclusions.

To estimate the overlap between the population activity in the two conditions, we calculated the alignment index. We slightly modified the calculation used in a previous study (Elsayed et al., 2016) to control for the variability that results from trial-by-trial fluctuations in activity. For each condition *i,* we defined *A_i_* as the activity matrix of all neurons and *PC_i_* as the matrix containing the first 5 PCs of the activity matrix *A_i_*. We denote *var*(*A_i_*·*PC_j_*) as the sum of the variances of the projection of activity in condition *i* on the PCs of condition *j*. We divided the trials of each neuron within each condition randomly into two groups, resulting in two different population activity matrices for each condition. We denote these matrices and the corresponding PCs as *A_i,k_*, *PC_i,k_* where k is the index of the group (1 or 2). This data division allowed us to control for the impact of noise on the PSTH estimation affecting the PCA, albeit with a minor increase in the noise.

The alignment index of j-th condition on the i-th condition was defined as:

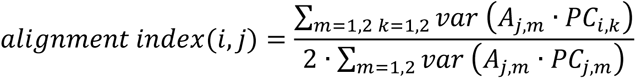

i.e., the variance of the projection of the activity in the j-th condition on the PCs calculated in the i-th condition normalized by the projection of the activity in the j-th condition on its own PCs. Note that the denominator is multiplied by 2 since it represents half the terms.

The alignment index of a condition on itself was defined as:

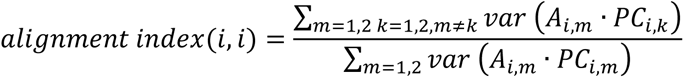

i.e., the variance of the projection of the activity in the two randomly divided groups on the other group’s PCs, normalized by the projection of each group’s activity on its own PCs. The alignment index of condition *i* on itself is negatively correlated with the trial-by-trial noise (minimal trial-by-trial noise will yield an alignment index close to 1), and provides a control for the size of the alignment index for conditions in which subspaces completely overlap.

We used the first 5 PCs that account for more than 55% of the variance for calculating the alignment index. We confirmed that using up to 10, that account for more than 70% of the variance, did not alter any of our conclusions.

To assess whether the alignment indices of the two different conditions to a third condition were significantly different, we conducted a permutation test. We randomly shuffled the condition labels of half of the neurons and used the difference in alignment index between the conditions as the test statistic. We repeated this permutation process 1000 times to create a distribution of the statistic under the assumption of no difference between conditions. The p-value was then defined as the probability of observing a more extreme difference in alignment index than the statistic for the permuted samples (two-tailed test). Note that this test assumes that the equality in alignment indices arises from the similarity in population responses between conditions, although it is possible that different population activity matrices could yield similar alignment indices.

We quantified the correlation of the directional tuning between tasks using the Pearson correlation. For each neuron, within each condition and for each 50 ms bin, we constructed a tuning curve. We calculated the correlation of the tuning curves between the conditions for each pair of conditions, in each bin where the neuron exhibited significant directional tuning in both conditions.

To calculate the population vector, for each neuron we constructed the tuning curve for the different tasks in the first 350 ms following motion onset. We then determined the preferred direction (PD) of the neuron in the pursuit task. To prevent bias, for each neuron, half of the trials were randomly selected to determine the PD and the other half to calculate the population vector. For a given condition and direction we defined the population vector on the horizontal and vertical axis as:

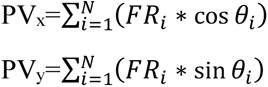

where FR_i_ is the firing rate of neuron *i* in the appropriate condition and direction, *θ_i_* the preferred direction of neuron *i* and N is the total number of neurons included in the analysis.

To compare the behavioral data with the population vector analysis, we utilized the behavioral data recorded while monitoring the neurons used for the population vector calculations. For each direction we calculated the average change in position in the horizontal and vertical axes between motion onset and 350 ms after motion onset across trials. Then, we averaged the changes in position along the horizontal and vertical axes across all neurons to determine the average change in position in the different conditions and directions.

To quantify the attenuation in neural activity between the suppression and pursuit conditions, we calculated the fraction of the root mean square of the population vector across directions as follows:

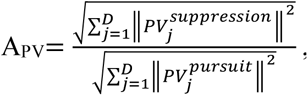

where *PV_j_^task^* is the population vector for a given task and direction *j* and || || is the Euclidean norm.

The attenuation in the change in position between suppression and pursuit conditions was calculated similarly.

## Results

### Possible organizations of population activity

The methods used to determine population organization are based on a comparison of the neuron response properties across tasks (e.g., directional tuning during different eye movements). Specifically, we examined whether the consistency in tuning between two neurons on one task persisted on another task. Figure 1 shows the possible set of relationships between the neuronal responses across tasks (Fig. 1, left and middle columns) as well as the geometric implications derived from the specific patterns of activity (Fig. 1, right column). For reference we consider two neurons that exhibited similar tuning on one task (Fig. 1A). Hence, plotting the activity of neuron 1 versus neuron 2 spans a specific subspace along the equality line (Fig. 1B).

**Figure 1:**
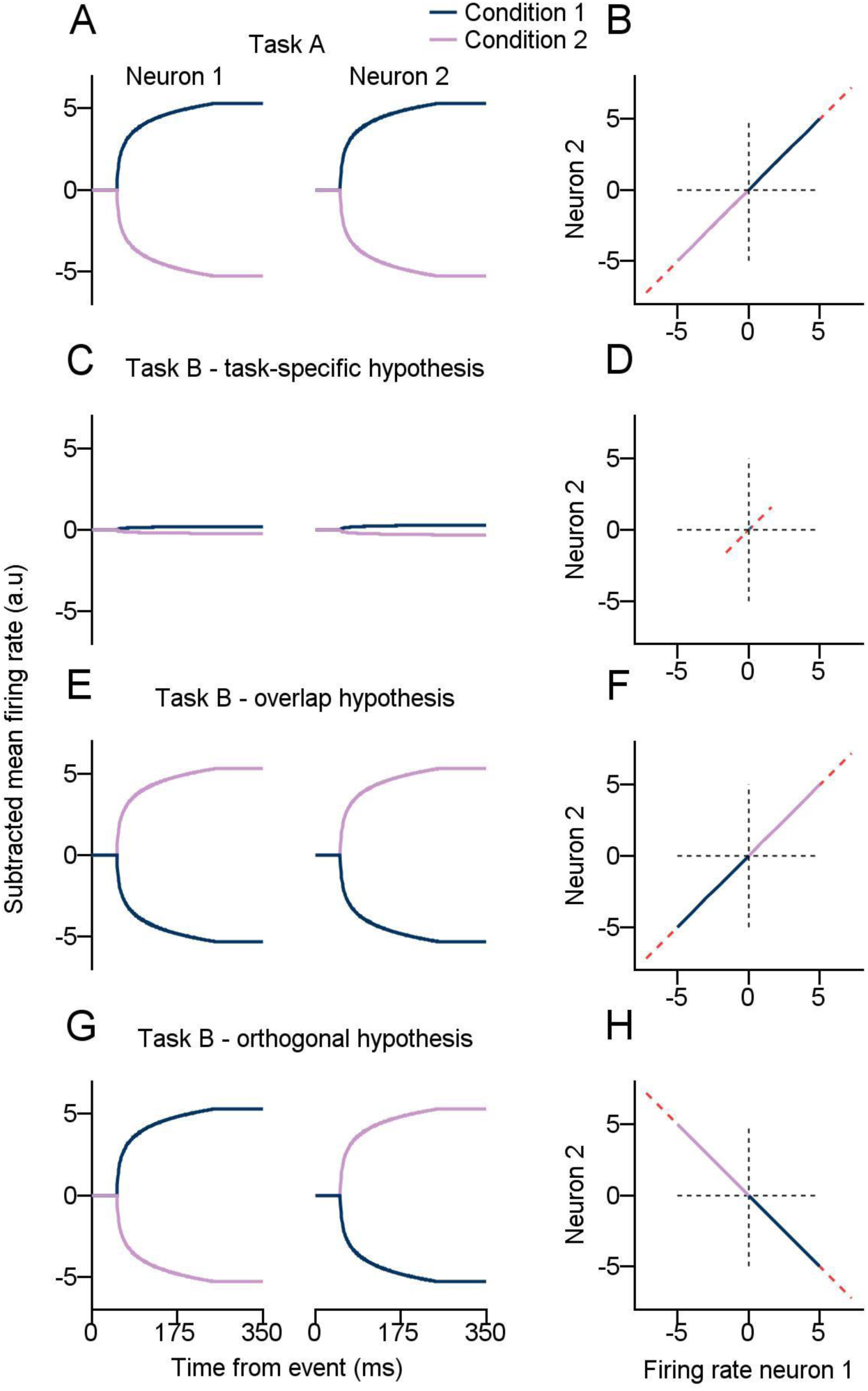
Illustration of hypothesized neural activity for a theoretical pair of neurons in two different eye movement tasks. **A:** Firing rates as a function of time for neurons 1 and 2 on task A. The color of the traces indicates the two task conditions (e.g., directions of movement). **C, E, G:** Firing rates as a function of time for neurons 1 and 2 on task B according to the three hypotheses: each task is coded by different neurons (**C**), the subspaces defined by the neural activity in different tasks overlap (**E**) and the subspaces defined by the neural activity in the different tasks are orthogonal (**G**). The colors of the traces indicate the task condition and correspond to the conditions in A (e.g., same movement direction). **B, D, F, H:** Firing rate of neuron 1 versus firing rate of neuron 2 on task A (**B**) and task B according to the three hypotheses (**D, F, H**). Neuronal activity defines a subspace (red dashed line) within the full space of potential states.

We explored three hypothetical strategies to distinguish between movements at the population level. The first was a task-specific hypothesis, where different clusters of neurons, within a specific area, encode different movement or sensory features (Fig 1C). For example, FEF neurons might exclusively respond to saccades or pursuit but not both. The second, the overlap hypothesis, posits that similar tuning on the first task coincides with similar tuning on another task. In this case, the activity will span the same subspace as observed in the first task (Fig. 1B versus Fig. 1F). For example, neurons in the final motor pathways of eye movements often consistently code the direction of movement across distinct eye movements (Sylvestre and Cullen, 1999). Note, however, that the consistency of neurons across tasks is sufficient but not necessary for a shared subspace, as shown by the switch in the preferred condition between Figure 1A and E. Finally, the orthogonal hypothesis posits that the neural activity on two different tasks presents inconsistencies. For instance, neurons may be similarly tuned in one task and oppositely tuned in the second task (Fig. 1G). In this case, activity on the two tasks will span orthogonal, non-overlapping subspaces (Fig. 1B versus Fig. 1H). This orthogonal pattern has been observed in the motor cortex, where population activity during the preparatory phase and actual movement execution displays no overlap (Elsayed et al., 2016).

In these examples of population-level organizations we used the simplified scenario of a population comprised of only two neurons. When we extend this concept to a higher-dimensional pattern involving numerous neurons, directly visualizing these subspaces becomes impractical. However, the same principles should hold at the population level: tuning consistency across tasks suggests overlap, whereas inconsistency implies orthogonality. The next sections discuss different eye movement tasks and present the analyses aimed at differentiating between these hypotheses.

### Tasks probing distinct categories of eye movements and the response of an example neuron

To investigate population coding across distinct categories of behavior, the monkeys were engaged in different eye movement tasks, while neural activity was recorded from the FEF. At the start of each trial, a white spot appeared in the center of the screen and the monkeys were required to fixate on that spot (Fig. 2A, left column). In the pursuit task, after 500 ms of fixation, an eccentric target appeared along one of eight possible directions to indicate the upcoming direction of motion (Fig. 2A, middle column, top). We defined this time point as the cue onset. After a variable delay, the eccentric target started moving towards and through the center of the screen at a speed of 20°/s and the monkey was required to track the target (Fig. 2A, B top). The eccentric position of the target served two purposes: it indicated the direction of motion and reduced the need for catch-up saccades at pursuit (similar to step-ramp motion; Rashbass, 1961). The suppression task was similar to the pursuit task, except that the central fixation target was a square that remained displayed during motion. The monkeys had to maintain fixation on the square central target to successfully complete the trial (Fig. 2A, B middle). Comparison of eye movement traces during pursuit and suppression (Fig. 2B, top versus middle) revealed that the monkeys effectively suppressed movement during the suppression task, although some residual movement still persisted. In the saccade task, the fixation target disappeared and reappeared at one of eight 10° eccentric locations and the monkeys were required to saccade towards this target (Fig. 2A, B bottom). No cue was displayed during the saccade trials. We defined motion onset in the pursuit and suppression tasks as the time point when the eccentric target started moving, and in the saccade task as the moment when the central target jumped.

**Figure 2:**
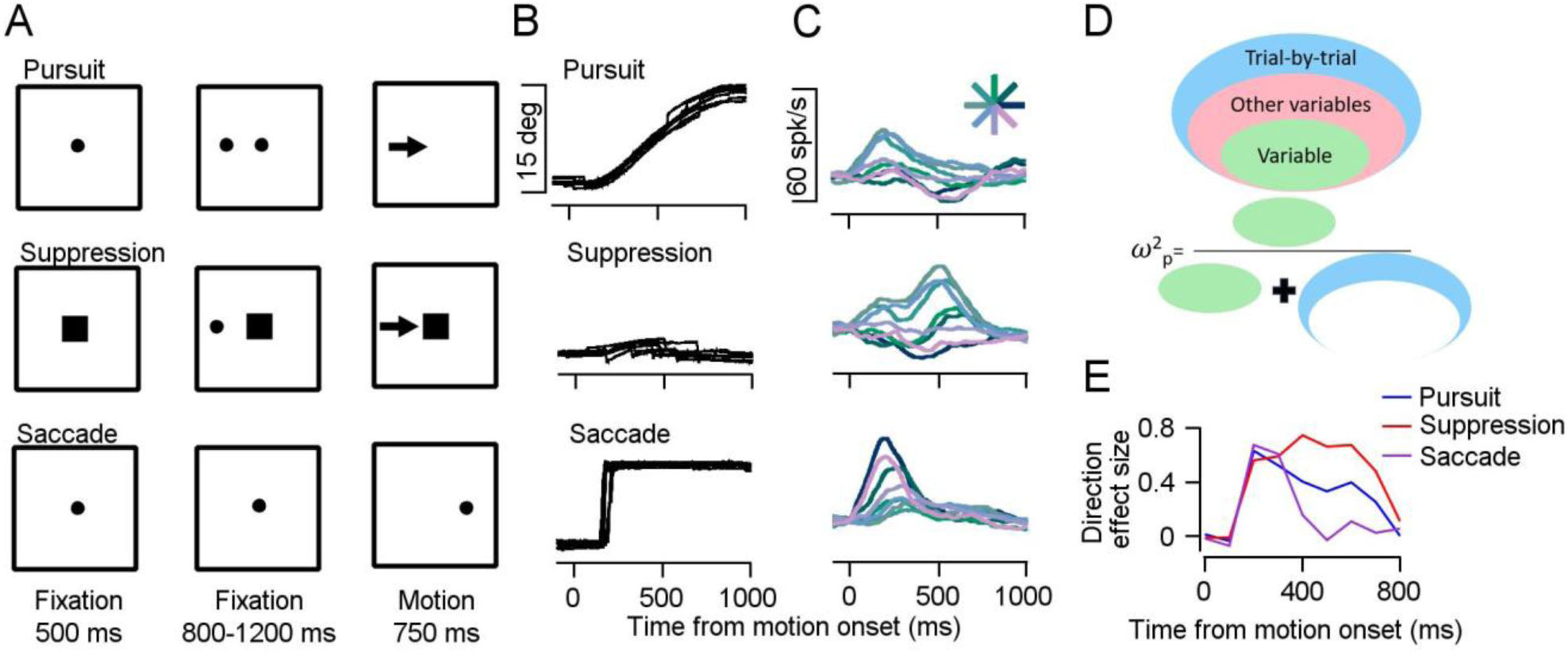
Behavioral task, example neuron and direction effect size analysis. **A:** Schematics of pursuit, suppression and saccade tasks as they appear on the screen. Spots and squares represent motionless targets and the arrows show the direction of continuous motion in the pursuit and suppression tasks **B:** Horizontal eye position aligned to motion onset for trials in which the target moved to the right from an example recording session. Each trace shows a different trial. **C:** PSTHs of an example neuron aligned to motion onset. Colors indicate the direction of target motion and correspond to the direction presented in the asterisk at the top of **C**. **D:** An illustration of the calculation of the *ω*^2^_p_ effect size. The ovals represent the partition of the trial-by-trial variability into the variance explained by a specific variable (blue), the variance explained by other variables (pink) and the variance unexplained by any of the task variables (green). **E:** Direction tuning effect sizes of the neuron shown in **C** calculated in bins of 100 ms.

While the monkeys performed these eye movement tasks, we recorded single-neuron responses from the FEF (497 and 716 neurons from monkeys F and J, table 1). To achieve representative sampling of the FEF, we used recording techniques that mitigate potential biases stemming from selective recording. We used multi-contact probes (Plexon probes with either 24 or 64 channels), which enabled us to record a multitude of neurons without pre-selecting the neuron based on the properties of the neural response.

**Table 1:** Number of neurons recorded. Overall column: number of neurons recorded in each task and combinations of tasks. Significant column: number of neurons with significant differences between activity in the first 350 ms after motion onset and the 400 ms before (Wilcoxon signed-rank test, critical value of 0.01).

| Row index |  | Monkey F |  | Monkey J |  |
| --- | --- | --- | --- | --- | --- |
|  |  | Overall | Significant | Overall | Significant |
| A | Saccade | 407 | 166 | 449 | 129 |
| B | Pursuit | 485 | 197 | 715 | 167 |
| C | Suppression | 485 | 188 | 715 | 108 |
| D | Cue (pursuit) | 485 | 83 | 715 | 70 |
| E | Saccade or Pursuit | 395 | 244 | 448 | 177 |
| F | Saccade or Suppression | 395 | 232 | 448 | 157 |
| G | Saccade or cue | 395 | 200 | 448 | 166 |
| H | Pursuit or Suppression | 485 | 244 | 715 | 195 |
| I | Pursuit or Cue | 485 | 235 | 715 | 208 |
| J | Suppression or Cue | 485 | 219 | 715 | 157 |
| K | Saccade and Pursuit | 395 | 87 | 448 | 70 |
| L | Saccade and Suppression | 395 | 88 | 448 | 50 |
| M | Saccade and cue | 395 | 40 | 448 | 20 |
| N | Pursuit and Suppression | 485 | 141 | 715 | 80 |
| O | Pursuit and Cue | 485 | 45 | 715 | 29 |
| P | Suppression and Cue | 485 | 52 | 715 | 21 |

The example neuron in Figure 2C illustrates the directional tuning during different tasks after motion onset (Fig. 2C). In particular, the tuning of this neuron at motion onset was similar for suppression and pursuit, as indicated by the similar order of colors on these tasks (Fig. 2C top and middle). On the saccade task, the directional tuning was different, as indicated by the different order of colors compared to the pursuit and suppression tasks. Thus, this neuron shows that neurons can respond with substantial modulation during pursuit, suppression and saccade movements. In addition, this neuron demonstrates that tuning across tasks may not be consistent. In the following sections we report analyses examining whether the same neurons responded across tasks and whether coding of motion direction was consistent across tasks at the level of single neurons and across the population. We first focused on the motion epoch of the three eye movement tasks, and leave the analysis of the cue epoch to later sections.

### FEF neurons are directionally tuned in different eye movement tasks

We first quantified the extent of directional tuning encoded on the different tasks for single neurons. To do so, we utilized the partial *ω*^2^ (*ω*^2^_p_, see methods) as a measure of the effect size of the directional tuning in the different conditions (Olejnik and Algina, 2000; Larry et al., 2022). *ω*^2^_p_ serves as an unbiased estimator of the variability explained by an experimental variable, divided by the sum of this same variability and a noise term (Fig. 2D, bottom). The noise term represents the variability that remains unaccounted for by any variable in the experiment, thus reflecting trial-by-trial variability within conditions. *ω*^2^_p_ values close to 1 indicate large differences between the levels of a variable relative to the noise within conditions, whereas values close to 0 indicate an absence of differences between conditions. Figure 2E shows the *ω*^2^_p_ calculated in bins of 100 ms for the example neuron. The increase in the values after motion onset indicates that the neuron exhibited clear directional tuning on the pursuit, suppression and saccade tasks.

We quantified the directional tuning in the FEF by averaging *ω*^2^_p_ across neurons (Fig. 3A). Pursuit, suppression and saccade tasks exhibited similar magnitudes in terms of direction effect size, indicating that the FEF was involved in encoding eye movement directions in all three tasks. The saccade tuning effect size was the largest around the time of the saccade but remained positive even after the saccade was completed (Fig. 3A, purple). On the pursuit and suppression tasks, the neuronal population continued to be strongly tuned throughout target motion and reached similar values (Fig. 3A, red and blue).

**Figure 3:**
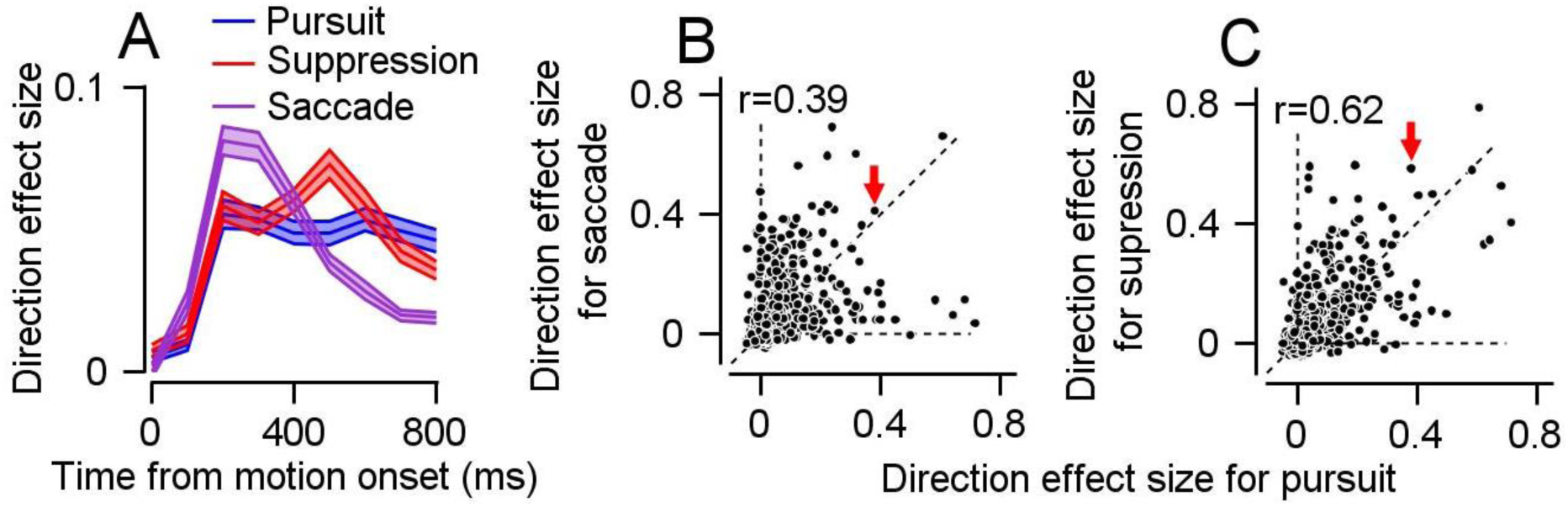
Direction effect size analysis shows that the same neurons are tuned on different tasks. **A:** Direction effect sizes averaged across neurons in bins of 100 ms. **B, C:** Comparison of the effect size on different tasks. Each dot represents a single neuron’s direction effect size on the pursuit (horizontal) and saccade (vertical) tasks (**B**) and the pursuit (horizontal) and suppression (vertical) direction effect sizes (**C**). The r values show the Pearson correlations. Dashed lines correspond to the identity line and cardinal axes. The red arrow points to the example neuron shown in Figure 1C.

### Single neurons are directionally tuned across distinct behaviors

One possible hypothesis concerning the ability to encode different movements is the division of the FEF into independent subpopulations responsible for different categories of eye movements (Fig. 1C). To investigate this hypothesis or whether many neurons exhibited directional tuning in more than one task, we calculated the effect size of each neuron during the 800 ms following the motion onset in each condition (see Methods). Many neurons had a positive directional effect size on more than one task (Fig. 3B and C) like the example neuron (marked by the red arrow in Figs. 3B and C). Overall, we found a substantial correlation in effect size between the pursuit and suppression tasks (r= 0.62), as well as between the pursuit and saccade tasks (r=0.39). The neurons were not distributed along the cardinal axes as expected from distinct coding of the different tasks, indicating that the same FEF neurons tended to be directionally tuned in different tasks rather than organized into distinct clusters for each eye movement task. Therefore, for the pursuit, suppression and saccades, our data contradicts the task-specific hypothesis (Fig. 1C) with respect to the organization of FEF population activity.

### The subspace defined by the population activity in the pursuit tasks overlaps with suppression and is mostly orthogonal to the saccade task

The lack of distinct clusters in the FEF led us to explore the organization of the FEF population during different eye movement tasks. Specifically, we tested whether the population activity in the different tasks overlapped (Fig. 1F) or would lie along orthogonal spaces (Fig.1H). Due to the intricate time dynamics of neurons in the FEF (see for example Fig. 2C, top), we decided to concentrate on the encoding of motion onset (0-350 ms). This focus also optimized the comparison between the pursuit and suppression tasks with the saccade task, where eye movements occur within a narrow time window after motion onset (Fig. 2B, bottom).

The key component of this population analysis was to test whether pairs of neurons consistently encoded responses across tasks (Fig. 1). The mathematical criterion to assess consistency in tuning between tasks is the similarity of the population covariance matrix on the two tasks. Thus, to study population organization, we first compared the covariation of neurons to test whether the neuronal tuning was consistent across tasks and then examined the structure of the covariance matrix by using a principal component analysis (PCA).

### Consistency of tuning across tasks

To assess whether pairs of neurons that shared a directional tuning pattern in one condition exhibited consistent response patterns on other tasks, we computed the covariance of the responses for each pair of neurons and eye movement task (see Methods). We refer to this covariance as the *tuning consistency score*. A positive tuning consistency score indicates that two neurons share consistent tuning on a given task, whereas a negative score indicates that the neurons exhibit opposite tuning patterns. We calculated the correlation coefficient of the tuning consistency score between pursuit and the two other tasks. This correlation quantified the extent to which the tuning consistency score was preserved between tasks. There was a significantly higher correlation in tuning consistency score between the pursuit and suppression tasks (r=0.56, Fig. 4A right) than between the pursuit and saccade tasks (r=0.07, Fig. 4A left, Fisher transformations, p<10^-3^, see Methods). This higher correlation manifested in the tendency of the scores for pursuit versus suppression tasks to distribute along the oblique direction whereas the distribution on the pursuit versus saccade tasks tended to be less structured (Fig. 4A). Thus, our results suggest that the subspaces occupied by the pursuit and suppression population responses substantially overlap (hypothesis presented in Fig. 1F), while the pursuit and saccade population responses exhibited more orthogonal subspaces (hypothesis presented in Fig. 1H). Although the correlation between tuning consistency scores is a relatively direct measure of consistency across trials, the values of the correlation should not be interpreted as the exact extent of the overall overlap.

**Figure 4:**
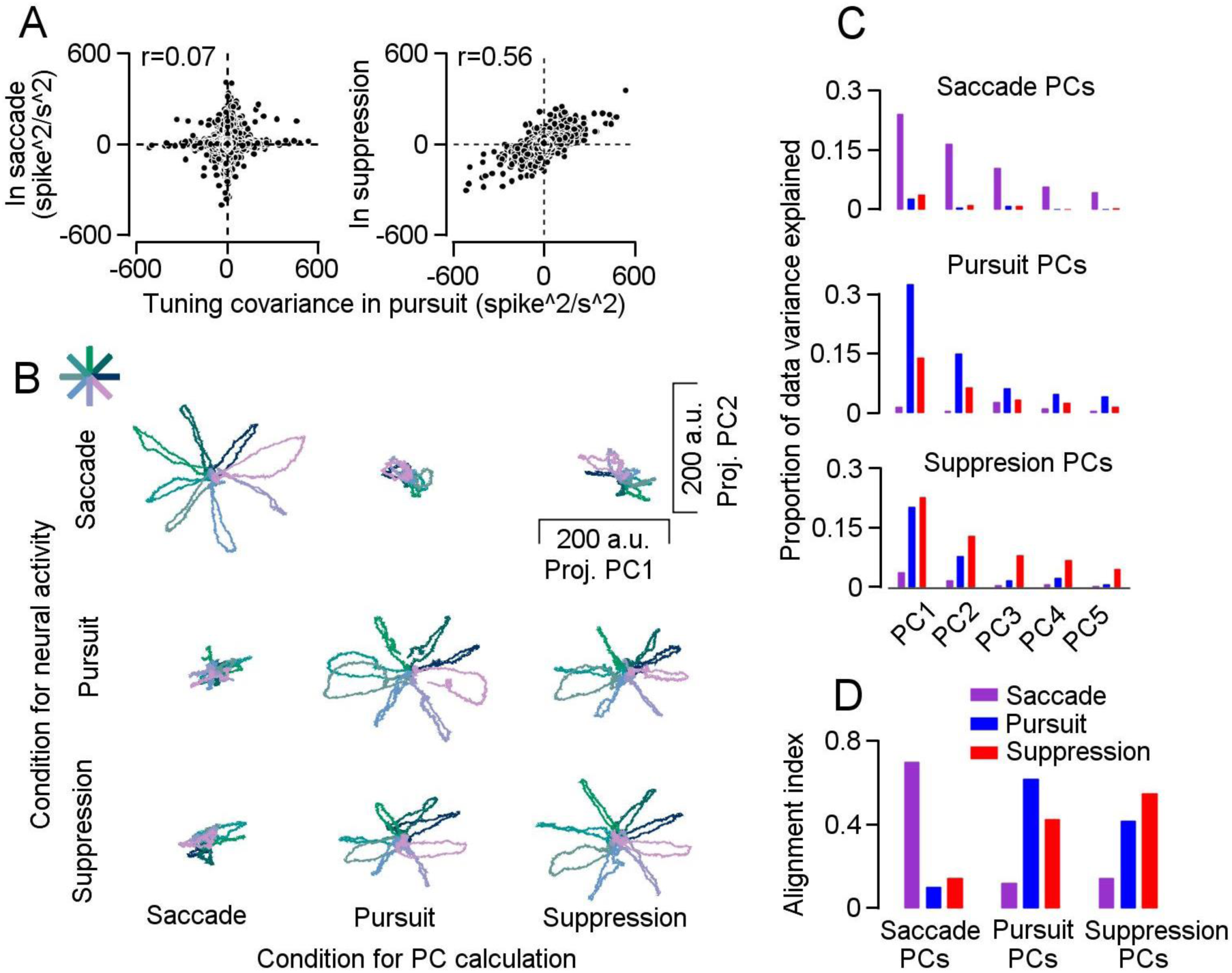
Population activity on the pursuit and suppression tasks share similar subspaces that differ from the subspace of activity during saccades. **A:** The covariance between responses (tuning consistency score) for all pairs of neurons on the pursuit task plotted against the covariance for the same pair during the saccade (left) and suppression (right) tasks. The r values represent the Pearson correlations. **B:** Projection of the population activity from one task onto the first two PCs calculated for the second task. Columns correspond to the task in which the PCs were calculated and rows correspond to the task used to project on the PC. Each colored trace plots the trajectory for one of the eight directions that correspond to the asterisk in the upper right panel. **C:** Proportion of variance on each task explained by the top five PCs for each task. **D:** Alignment index for all possible combinations of pairs of tasks on the subspace defined by top-five saccade PCs (left), pursuit PCs (middle) and suppression PCs (right).

### Visualization of the overlap between subspaces with leading principal components

To assess the overlap between population activity on the three tasks we used dimensionality reduction methods. For each task, we ran a principal component analysis (PCA) where the principal components (PCs) were defined as the eigenvectors of the covariance matrix (see Methods). In Figure 4B, for each task, the diagonal graphs show the projection of the task population response on the first and second principal components which were the two directions that explained the largest percentage of the variance (38%, 47% and 41% on the pursuit, suppression and saccade tasks). As expected from the large tuning effect size of many neurons (Fig. 3A and B), there were clear, different patterns for each direction (diagonal plots in Fig. 4B). In off-diagonal plots, we plotted the projection of the population response for each task on the PCs of the other tasks. The pattern of activity of these projections enabled us to visually compare the overlap in the subspaces of the activity across the three tasks in the lower-dimensional space defined by PC1 and PC2. In line with the high correlations in the tuning consistency scores, projecting the pursuit activity on the suppression PCs (and vice-versa) preserved the structure of the data (lower-rights plots in Figure 4B), supporting the hypothesis of an overlap of the subspaces spanned by the pursuit and suppression population responses. Conversely, projecting the saccade population response on the pursuit or suppression PCs (and vice-versa) mostly collapsed the structure of the data, thus corroborating the hypothesis of orthogonal spaces between saccade and pursuit or suppression population activity.

We then expanded our analysis beyond the first two principal components by projecting the population activity onto additional principal components and calculating the proportion of variance within the total dataset variance accounted for by each PC (Fig. 4C, see Methods, first 5 PCs accounted for 64%, 57% and 62% of the variance on the pursuit, suppression and saccade tasks). Projection of the population activity during saccade task on the pursuit and suppression PCs (and vice-versa) resulted in small values, indicating that these PCs only accounted for a small portion of the total variance in saccade activity (blue and red bars at the top of Fig. 4C and purple bars in lower plots of Fig. 4C). By contrast, the projection of pursuit activity on the suppression PCs (and vice-versa) yielded much larger values, indicating that the leading PCs in the suppression task explained a significant portion of the variance in the pursuit activity (red bars in the middle of Fig. 4C and blue bars in the bottom of Fig. 4C).

### Quantifying the overlap between subspaces

Next, to quantify the overall overlap between subspaces, we calculated the alignment index (Elsayed et al., 2016) (see Methods). This index assesses how much of the variability is preserved when we project the population activity from one task onto the PCs computed in another task. If the subspaces spanned by the population activity across tasks are orthogonal (no overlap), the index will be close to 0. Conversely, a theoretical maximum value of 1 suggests complete overlap between the subspaces. However, practical constraints, such as trial-by-trial fluctuations prevent reaching a value of 1 even if two subspaces completely overlap. To make the alignment index more interpretable, we employed a method where we computed the PCs using half of the trials and then used the remaining half to project the data (see Methods). We used the projection of each task onto itself to establish a practical measure of the maximal overlap between spaces (the highest purple, blue and red bars in Fig. 4D).

Figure 4D shows the alignment index for all pairs of tasks, and depicts the larger overlap between the pursuit and suppression subspaces than for the saccade. For example, the alignment index computed by projecting the suppression population response onto the space defined by the pursuit PCs (0.42) significantly exceeded that of projecting the saccade population response onto the pursuit PCs (0.13, Fig. 4D, permutation test, p<10^-3^). The overlap between the suppression population response and the pursuit PCs was 68% of the alignment index of the pursuit activity with itself (0.62, see Methods). Overall, and consistent with the overlap hypothesis (Fig. 1F), we found a substantial overlap between the subspaces defined by the pursuit and suppression population activity. Conversely, consistent with the orthogonal hypothesis (Figure 1H), the subspaces spanned by the population response in saccade and pursuit (or suppression) were mostly orthogonal.

### Pursuit and suppression tasks exhibit similar encoding in the FEF

So far, we have shown that the high correlation of the tuning consistency score between pursuit and suppression tasks was reflected in the substantial overlap between the subspaces spanned by the population responses on these tasks. Recall that this correlation indicates that neurons with similar tuning on the pursuit task are likely to exhibit similar tuning on the suppression task. However, this finding does not necessarily imply that the tuning of each neuron will be similar between tasks. For instance, a pair of neurons with preferred directions (PD) of 0° on the pursuit task and 180° in the suppression task (e.g., Fig. 1A and E) would still yield a high tuning consistency score. This has important functional implications, since differences in tuning are frequently utilized by commonly assumed decoders such as population vectors (Georgopoulos et al., 1983) to generate differential output from FEF activity. Conversely, similarity in tuning would imply that the population vector cannot distinguish between pursuit and suppression, thus necessitating additional mechanisms to account for the behavioral readout during pursuit and suppression. For these reasons, we next focused on the consistency of directional tuning at the individual neuron level across tasks.

To study the extent to which directional tuning was preserved between tasks, we first calculated the average population PSTH across neurons for all directions relative to the preferred direction (PD) on the pursuit or suppression tasks (Fig. 5A, diagonal). Next, for each neuron we ordered the PSTH based on the PD in the other task (Fig. 5A, off-diagonal). Calculating the population PSTH based on the pursuit or suppression task did not markedly change the population tuning (Fig. 5A left, top versus bottom for pursuit aligned with suppression and Fig. 5A right for suppression aligned with pursuit). This finding suggests that directional tuning remains very similar between the suppression and pursuit tasks after motion onset.

**Figure 5:**
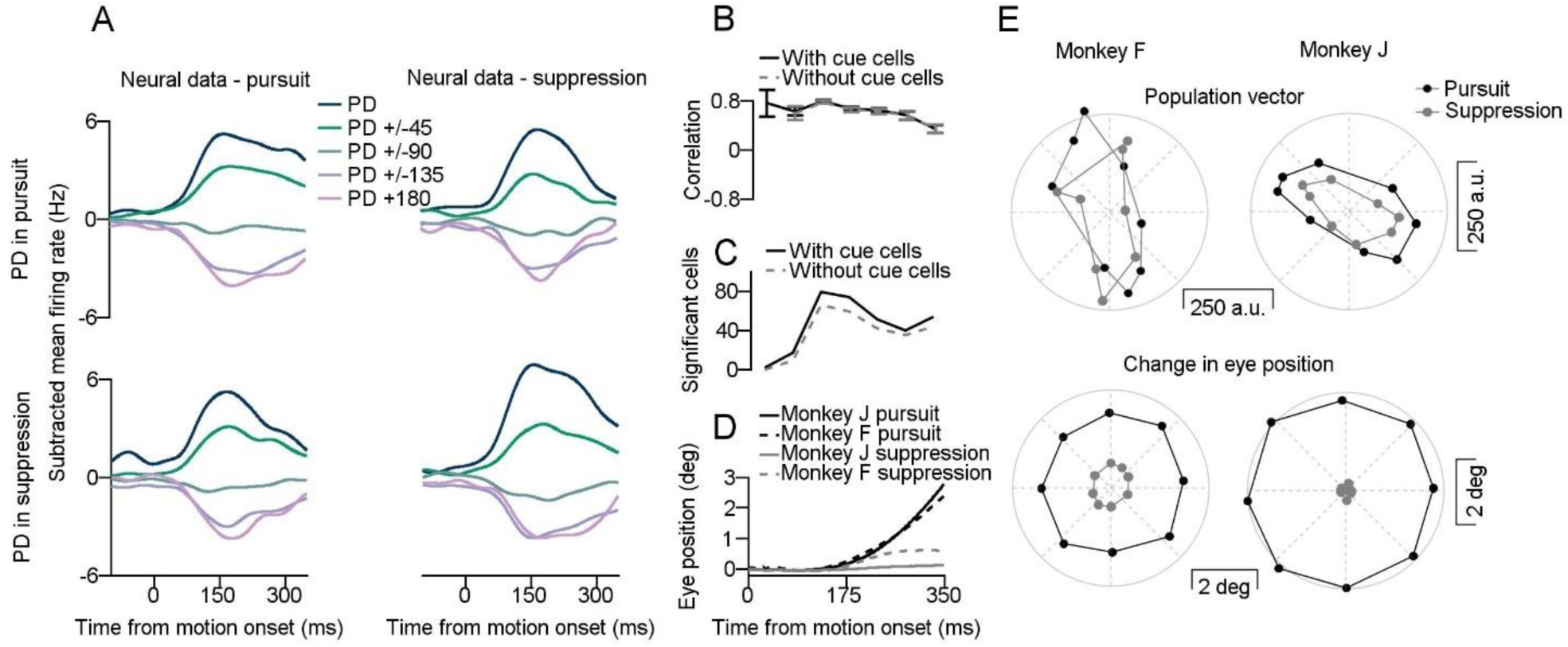
Similar population averages and directional tuning in pursuit and suppression tasks. **A:** Population average of the neural activity for the pursuit (left column) and suppression (right column) tasks. Different plots correspond to the differences in the direction of motion from the PD of the neurons computed for the pursuit (upper row) and suppression tasks (bottom row). The mean firing rate was defined as the average rate at motion onset across directions and was subtracted individually for each neuron. **B:** Average correlation across neurons in directional tuning between the pursuit and suppression tasks in 50 ms time bins. Solid and dashed lines show results that include and exclude neurons significant for directional tuning in the 400 ms before motion onset. Only neurons with significant directional tuning in both conditions in the relevant bin were included in the analysis. **C**: Number of neurons with significant directional tuning in the relevant bin. **D:** Eye position aligned to motion onset for monkey F (dashed lines) and J (solid lines) in pursuit (black lines) and suppression (grey lines) tasks **E:** Top: Population vectors based on the neurons’ PD on the pursuit task calculated for the pursuit and suppression tasks for monkey F (left) and monkey J (right). Each dot represents the population vector readout for one direction of motion. Bottom: Average change in eye position during the pursuit and suppression tasks for monkey F (left) and monkey J (right). Each dot represents the eye displacement for one direction of motion.

To quantify this similarity, we calculated the average correlation across neurons in directional tuning between the pursuit and suppression tasks in 50 ms time bins (Fig. 5B). This analysis only included neurons that showed significant directional tuning in both tasks within the relevant bin (see Methods, Fig. 5C). The results revealed a high correlation in tuning between tasks, especially immediately after motion onset, indicating that neurons in the FEF exhibited similar tuning on the pursuit and suppression tasks after motion onset. To ensure that this correlation was not influenced by the processing of the cue which was identical in both tasks, we excluded neurons that were significantly tuned before motion onset. This exclusion did not affect the results (Fig. 5B, C gray dashed line). Thus, the results point to a considerable degree of similarity in the response of single neurons between the pursuit and suppression tasks.

To study the implications of similarity in tuning and magnitude of the responses, we used the population vector as a readout of the activity from the FEF (see Methods). The analysis of the population vector complements the analysis of the principal components in the sense that it projects activity on two specific directions in the neural space which are a-priori thought to be important. In contrast, the PCA selects directions based on the actual variance in the data. Note, however, that the analysis is not independent since the leading directions found by the PCA could (and do) overlap with the direction on which activity is projected in the population vector.

To compare the population vector during pursuit and suppression we calculated the population vector on both tasks based on the preferred directions of the neurons in pursuit. In this analysis we split the data into training and testing sets to obtain an unbiased estimation of the population vector (see Methods). The findings showed that the suppression population vector was very similar to the pursuit population vector (Fig. 5E top). For each condition, we calculated the average magnitude across population vectors for a given condition by calculating the root mean square (RMS) of the population vectors across directions (see Methods). The ratio of the average magnitudes of the population vectors in the pursuit and suppression conditions was 70% for monkey J and 90% for monkey F.

Importantly, the population vector readout could be compared to the actual eye movement readout to test whether similarity in activity arose from behavioral parallels between the tasks. Despite the fact that the monkeys were required to maintain fixation on the center of the screen during the suppression trials, minor residual eye movements were still present (Fig. 2B). Nonetheless, the attenuation in behavior was much stronger than the attenuation in neural activity. To demonstrate this point, we plotted the horizontal and vertical displacement in eye position in the first 350 ms after motion onset in the pursuit versus the suppression trials. The amplitude of the position displacement during suppression was 0.3 and 0.06 of the amplitude during pursuit for monkeys F and J (Fig. 5E bottom). Together, these findings provide further support for the hypothesis that there may be a comparable encoding mechanism between pursuit and suppression tasks. This raises the issue of the site of this divergence between tasks within the eye movement pathway, given the distinct behaviors they elicit (see Discussion).

### Pursuit and saccade exhibit weak single neuron tuning correlation between tasks

We found that activity during saccade and pursuit predominately occupied separate subspaces (Fig. 4). However, a closer examination of the projection of saccade on pursuit or suppression subspaces indicated some structure in the residual (Fig. 4B). Consequently, our next step was to study the relationship between tuning during saccade and pursuit tasks directly, by focusing on the subset of neurons tuned in both tasks. Specifically, we examined the consistency of the directional tuning between the two tasks at the single neuron level. Interestingly, we identified a reversal in directional tuning between these tasks, as illustrated by the reverse order of colors in the diagonal compared to the off-diagonal plots in Figure 6A. This reversal was further substantiated by the negative average correlation in tuning between the pursuit and saccade tasks shortly after motion onset (Fig. 6B). Thus, decoders relying on directional tuning would predict opposite readouts in these two tasks. This observation contrasts with actual eye movements, where direction is preserved between corresponding conditions (Fig. 2B). These findings further emphasize the distinctions between activity in the FEF and the readout of behavior. We revisit this point in the discussion.

**Figure 6:**
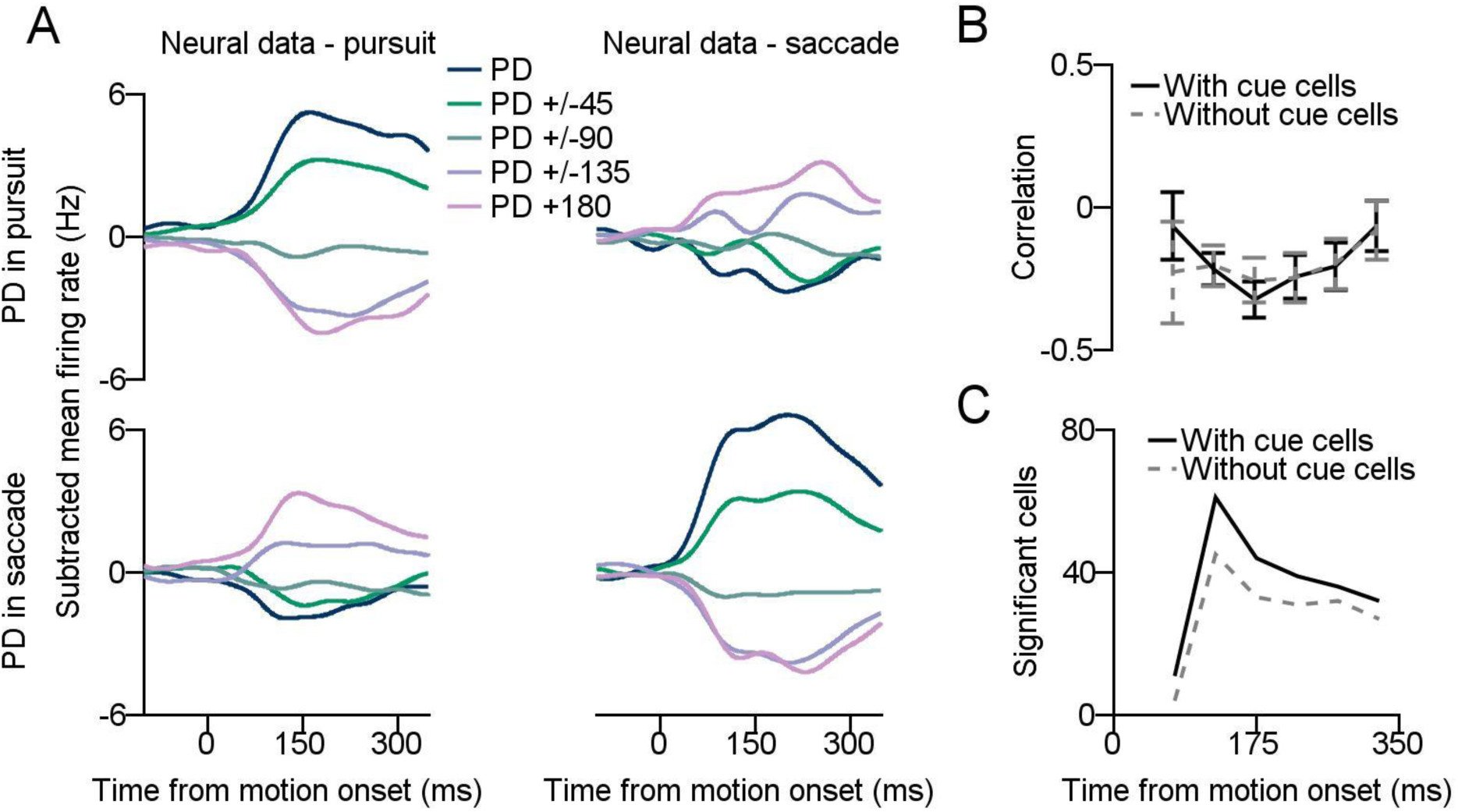
Weak tuning correlation between the saccade and pursuit tasks. **A:** Population average of the neural activity for the pursuit (left column) and saccade (right column) tasks. Different plots correspond to the differences in the direction of motion from the PD of the neurons computed for the pursuit (upper row) and saccade tasks (bottom row). The mean firing rate was defined as the average rate at motion onset across directions and was subtracted individually for each neuron. **B:** Average correlation across neurons in directional tuning between the pursuit and saccade tasks in 50 ms time bins. Solid and dashed lines show results that include and exclude neurons significant for directional tuning in the 400 ms before motion onset. Only neurons with significant directional tuning in both conditions in the relevant bin were included in the analysis. **C**: Number of neurons with significant directional tuning in the relevant bin.

Furthermore, we noted a significant decrease in neural activity during the saccade task when the PD of the neurons was determined according to the pursuit task activity (Fig. 6A). The root mean square (RMS) of the concatenated PSTHs in the saccade condition with the PD determined in the pursuit task was only 43% of the RMS of the concatenated PSTHs when the PD was determined based on the saccade task activity. In comparison, the suppression activity showed a less pronounced attenuation, with only 72% reduction (Fig. 5A). Note that these values do not imply a significant overlap between saccade and pursuit population activity since this analysis only considered neurons that modulated their response after motion onset compared to baseline in both tasks (19%, see Table 1, Methods). In fact, neurons modulated in only one of the two tasks could contribute to the analysis of the overlap but not to the PD and correlation analysis.

The quantification of correlations between tuning curves provided further support for the predominantly independent tuning of saccades and pursuit. The correlation between pursuit and saccades tasks was negative with a lower magnitude than the correlation between pursuit and suppression (Fig. 5B, C versus Fig. 6B, C). This difference in magnitude did not stem from directional tuning during the end of the cue epoch (dashed lines in Fig. 6B, C). Taken together, these findings provide further evidence for a disparity in the coding of pursuit and saccade based on the activity across tasks at the single-neuron level.

### Activity following cue onset is similar in pursuit and suppression trials and smaller than after motion onset

So far, our focus has been on analyzing activity during a range of behaviors after the initiation of motion. We now shift look at how the cue epoch was represented in the pursuit and suppression tasks (Fig. 7A). We assessed the magnitude of the directional tuning effect after cue onset in pursuit and suppression trials by averaging the values of *ω*^2^_p_ across neurons. The effect size during the cue onset was notably smaller than the effect size after motion onset and positive for a shorter duration compared to the effect size during motion onset (Fig. 7B top). To further study the distribution across neurons, we plotted the direction effect size in the different conditions as a function of the effect size in the cue pursuit condition (Fig. 7C upper right and bottom). Overall, directional tuning was more prominent after motion onset than after cue onset, as indicated by the bias towards the vertical axis in Figure 7C (p <10^-30^, for all tasks, Wilcoxon signed-rank test). In addition, a large proportion of the neurons was only tuned in the motion epoch, as indicated by the large number of data points along the vertical axis in Figure 7C.

**Figure 7:**
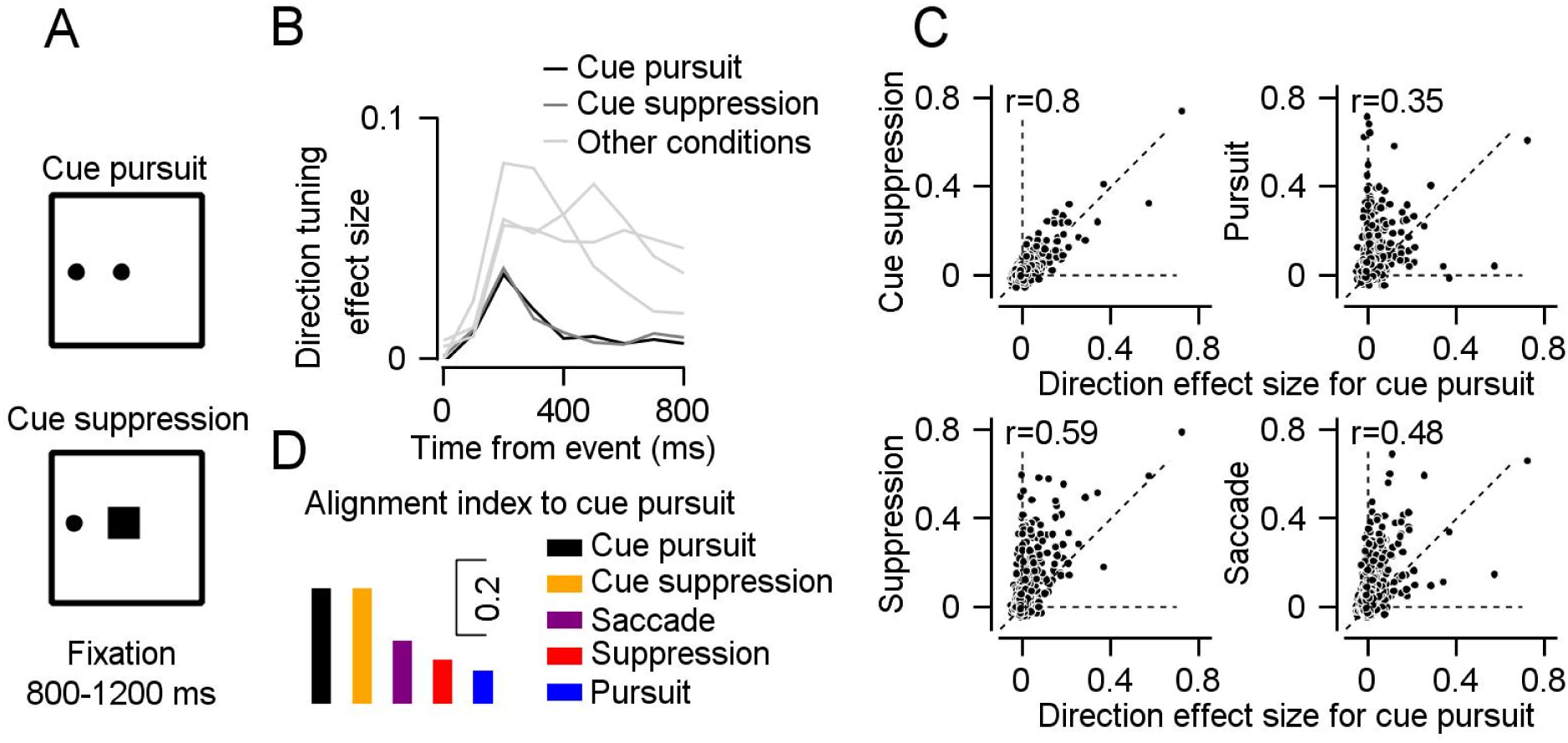
Cue-pursuit and cue-suppression exhibit similar responses. **A:** Schematics of pursuit and suppression as they appeared on the screen during the cue period. Spots and squares represent motionless targets. **B:** Average directional tuning effect sizes averaged across neurons in bins of 100 ms **C**: Comparison of the directional tuning effect size on different tasks. Each dot represents a single neuron’s directional effect size for each condition. Horizonal axis shows the effect size for the cue pursuit and compared along the vertical axis to cue suppression (**top-left)**, pursuit (**top right),** saccade (**bottom right)** and suppression (**bottom right**). The r values show the Pearson correlations. Dashed lines correspond to the identity line and cardinal axes **D:** Alignment index to cue pursuit PCs.

We did not find evidence for coding of the cue context (suppression or pursuit task). The direction effect size was similar in both conditions (Fig. 7B) and strongly correlated across neurons (Fig. 7C upper left panel, Pearson correlation=0.8). Furthermore, only a minimal percentage of neurons, (5.8%, 69 out of 1200, 2-way ANOVA, with direction and context), showed significant differences in responses between the two contexts. In addition, 4.5% of the neurons (54 out of 1200 neurons, 2-way ANOVA, with direction and context) showed a significant interaction between context and direction. This percentage did not deviate significantly from what would be expected by a chance at .05 (p=0.24 and p=0.44 for the main effect of context and the interaction between context and direction, χ^2^ test). For comparison 22% of neurons (260) exhibited significantly different responses between the pursuit and suppression tasks in the 350 ms following motion onset, which was notably different from what would be expected by chance (p<10^-5^, χ^2^ test) and larger than the fraction of neuron that responded significantly to the cue context (p <10^-5^, χ^2^ test).

The similarity in responses to the pursuit and cue suppression suggests that activity in the FEF before movement was unrelated to movement preparation. The lack of specific responses to context is consistent with a previous study that found limited modulation associated with proactive suppression in a pursuit GO/No-Go task (Fukushima et al., 2011). As expected from this similarity of responses, we confirmed that at the population level, the subspaces for cue pursuit and cue suppression overlapped highly (alignment index = 0.31 which was similar to the alignment index of each condition with itself - see Methods). The alignment indices of the other conditions to the cue-pursuit PCs were significantly lower and were overlapped to some extent (0.17 for saccade, 0.12 for suppression and 0.09 for pursuit, Fig. 7D).

### The FEF does not exhibit task-specific clustering at the anatomical level

In the previous section, we challenged the task-specific hypothesis (Fig. 1C) concerning the organization of FEF population activity. We did so by showing that individual neurons within the FEF exhibit directional tuning across various tasks, rather than being organized into distinct functional clusters for different eye movement tasks. This lack of functional specificity also implies a limited potential for specific anatomical organization. To test whether the absence of task-specific clusters at the population level was also evident at the anatomical level directly, we examined each recording location to determine whether at least one neuron exhibited significant tuning in different conditions (Kruskal-Wallis test). Overall, there were no clear anatomical task-specific clusters, although we did identify certain areas that appeared to be more closely associated with specific tasks. For instance, in the case of monkey J, on the posterior slice (AC+2), we identified a hotspot that was specific to pursuit. In summary, the absence of task-specific anatomical clusters supports the notion that the FEF’s ability to encode categorically distinct movements is rooted in the task-dependent organization of the neuronal population rather than in its anatomical structure.

## Discussion

The results showed that individual FEF neurons responded to different eye movement tasks (Fig. 3A) and displayed no task-specific clusters (Fig. 3B and C). We then identified a context-dependent organization at the population level. We observed a substantial overlap in the subspaces defined by the population response during the pursuit and suppression tasks (Fig. 4). By contrast, we found that the subspaces defined by the population responses on the saccade and pursuit tasks were mostly orthogonal (Fig. 4). These findings suggest that the subspace governing population activity depends on the context, as characterized by the type of motion of the moving target (smooth motion or ballistic jump), rather than on different categories of behaviors. In the following sections we discuss the implications of these results for the encoding of behavior in the FEF and the control of behavior across the eye movement system.

### Movement is suppressed downstream to the FEF

The substantial overlap in the subspaces defined by the population response in the pursuit and suppression tasks, along with the high positive correlation in directional tuning at the single-neuron level between these tasks (Fig. 4D and 5B) indicates that pursuit and suppression tasks are encoded similarly within the FEF. The substantial attenuations in behavior across tasks, compared to the similarity of encoding and readout (Fig. 5E), suggest that FEF neurons on the pursuit and suppression tasks are more tightly related to the motion of the target rather than to the movements of the eye. This dissociation between neural activity and behavior suggests that the decision to inhibit movement involves additional inhibitory network processing, either in parallel or downstream to the FEF. These cortical and subcortical networks may include the pontine omnipause neurons in the brainstem (Büttner-Ennever et al., 1988), cerebellar neurons in the vermis (Kurkin et al., 2014) or fixation neurons in the frontal cortex (Izawa et al., 2009).

One proposed mechanism for movement suppression involves the orthogonalization of the population response. For example, in the motor cortex, activity before and during movement exhibits orthogonal patterns (Elsayed et al., 2016). This orthogonality enables linear readouts that trigger movement exclusively during the movement phase (Kaufman et al., 2014). However, our results indicate that the absence of movement in the suppression task was not achieved through orthogonalization of the population response. Rather, the overlap between the population activity in the pursuit and suppression tasks implies that linear readouts of the pursuit activity, relying on dimensions containing substantial portion of the variability in FEF activity, would evoke movement during suppression.

For similar reasons, the partial overlap between the population activity during the cue and the saccade epochs (Fig. 7D) indicates that during the cue epoch, the suppression of a saccade towards the eccentric cue was not achieved only through an orthogonal organization of population activity. In addition, this suppression is not achieved through highly specific anatomical connections of neurons with motor-only responses (Segraves and Goldberg, 1987), indicating that further mechanisms are needed to prevent saccades, such as threshold crossing (Hanes and Schall, 1996) or downstream inhibition mechanisms (Büttner-Ennever et al., 1988). Yet, although orthogonalization is not the sole mechanism for the suppression of eye movement, the differences in the saccade and cue subspaces could contribute to preventing saccades during the cue.

### Implications of the common encoding of pursuit and suppression tasks on gain control by the FEF

The similarity between pursuit and suppression tasks challenges the hypothesis that the FEF primarily plays a central role in setting the gain of pursuit (i.e., the ratio of eye to target velocity) rather than directly encoding target velocity (Tanaka and Lisberger, 2001; Lee et al., 2013). According to the gain hypothesis, FEF activity should have decreased during suppression compared to pursuit. However, one possible explanation for this disparity is that the gain hypothesis has mainly examined FEF activity with moving targets, unlike the fixation targets used in this experiment (Lee et al., 2013; Darlington and Lisberger, 2020). Hence, it is possible that an active fixation mechanism (Raybourn and Keller, 1977) suppresses movement downstream from the FEF while the increase in activity during suppression enhances internal processing. This internal enhancement could facilitate the processing of visual information, which would be consistent with the motor origin of attention hypothesis (Moore et al., 2003). While most gain studies have involved moving targets, electrical stimulation in the FEF during fixation was shown to increase pursuit gain (Tanaka and Lisberger, 2001), thus further research is needed to reconcile this result with our findings.

### Implications of the orthogonality between pursuit and saccade subspaces

We corroborated the findings of a recent study (Mayo, 2022) that documented directional tuning in the same FEF neurons during pursuit and saccades (Fig. 3B). Initial studies identified the pursuit area within the FEF based on techniques such as electrical stimulation, neural activity recordings, and lesions (Macavoy et al., 1991; Gottlieb et al., 1993, 1994). However, these earlier investigations identified neurons involved in pursuit without systematically characterizing their responses during saccades to assess task specificity. In the current study, although we identified some neurons specific to the pursuit task (Table 1), we found that overall, the effect size of directional tuning between pursuit and saccade tasks was correlated (Fig. 3B) indicating that a significant proportion of the neurons responded to both tasks. In fact, even in the early studies, neurons that responded to both pursuit and saccades were noted (Tanaka and Fukushima, 1998). Furthermore, the sporadic nature of task-specific neurons is reflected in the absence of clear anatomic, task-specific clusters (Fig. 8), suggesting a more intricate organizational structure (Gottlieb et al., 1994).

**Figure 8:**
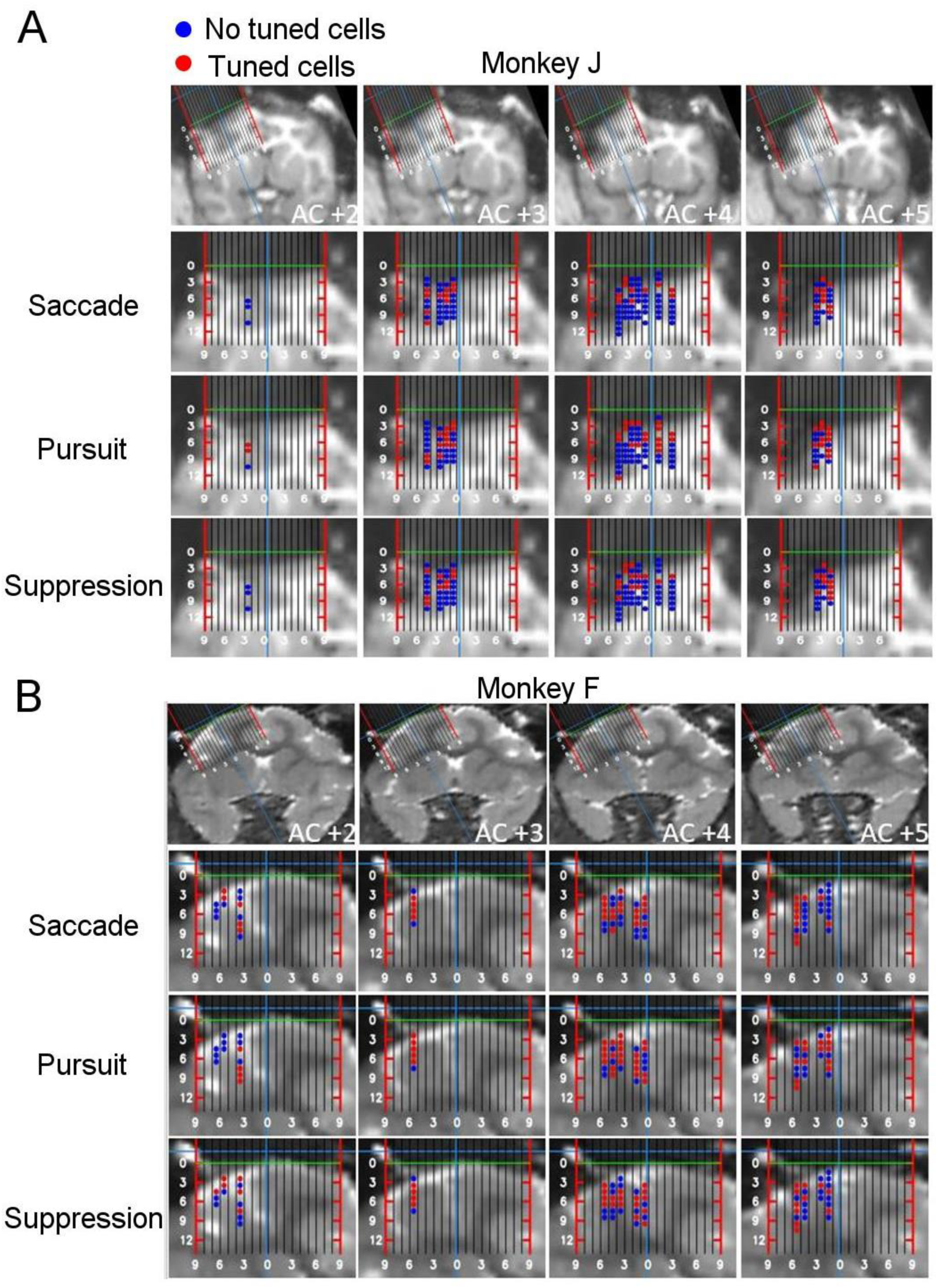
Anatomical organization of the responses in the FEF. **A, B:** The top row shows coronal MRI images acquired from monkeys J (**A,** T1 scan) and F (**B,** T2 scan), annotated with respect to the anterior commissure (AC). MRI images of monkeys J are horizontally flipped for consistency. The MRIs also incorporate a representation of the coordinate system employed during the recording in mm. The lateral edges of the coordinate system are depicted by the red bars. Green line represents the plane in which we first identified the cortex. The subsequent rows depict whether the recorded site included neurons that displayed significant tuning in the saccade, pursuit and suppression tasks for the corresponding coronal slice, overlaid on an enlarged view of the area marked by the coordinate system.

Rather than an anatomical or functional specificity in the responses, we identified an organization at the population level. The subspaces defined by the population activity in pursuit and saccade tasks were mostly orthogonal (Fig. 4D, left columns). Both subspaces led to movement, but each subspace encoded a distinct type of movement. This contrast with coding of multiple behaviors within overlapping subspaces in the motor cortex (Gallego et al., 2018). The orthogonality between the pursuit and saccade subspaces suggests that the “saccade-output-potent” subspace defined by the population activity in the saccade task lies within the “pursuit-output-null” subspace of the pursuit task and vice-versa. This orthogonal structure indicates that the readout of the FEF using two different linear decoders, each utilizing subspaces that capture most of the variability, may distinguish between the tasks.

Despite the predominantly orthogonal nature of the pursuit and saccade subspaces, a small amount of overlap persisted. On average, the FEF neurons exhibited slight negative directional tuning correlation between tasks. This negative correlation was evident in the reversal of directional tuning between tasks (Fig. 6A and B) and can be attributed to different target positions at motion onset. Specifically, the step-ramp-like paradigm used in the pursuit trials positioned the target at motion onset in the opposite direction to the saccade trials with the same movement direction. For example, in rightwards trials, the target location at motion onset in the pursuit trials was to the left of the screen center, while in the saccade trials, it jumped to the right. Neurons encoding target position with a PD to the right may have increased their firing rate relative to baseline on the saccade trials while decreasing this rate on the pursuit trials. This difference in target location relative to the screen center between saccade and pursuit tasks, combined with the presence of FEF neurons encoding target locations, may account for this negative correlation. Note , however, that only a small fraction of FEF activity could be attributed to the visual response since the cue response was very small compared to other events (Fig. 7B) (Goldberg and Bruce, 1985). This indicates that although the FEF might encode visual parameters, it is not coding visual information per se.

## Conclusion and future directions

The findings suggest that the FEF’s ability to encode both saccadic and pursuit eye movements stems from its population-level organization. These findings extend previous studies that have concentrated on individual neuron-level analyses, and point to the value of adopting a system-level perspective when investigating motor systems. The results contribute to a better understanding of the FEF organization. An intriguing direction for future research could involve examination how the population coding of the sensorimotor parameters we have identified interacts with established coding in the FEF of reward (Ding and Hikosaka, 2006; Lixenberg and Joshua, 2018), memory (Bruce and Goldberg, 1985), attention (Moore and Zirnsak, 2017) and learning (Li and Lisberger, 2011). In addition, we showed that pursuit and suppression tasks are encoded in a similar manner in the FEF. Future research could compare the activity in these tasks in areas downstream from the FEF within the eye movement pathway to investigate where and how the differentiation between the tasks occurs. Finally, the orthogonality of the saccade and pursuit subspaces indicates that readout could separate saccade and pursuit activities. This raises the question of whether communication between FEF and its output areas utilizes readouts that differentiate between saccade and pursuit activity.

## Acknowledgments

This project received funding from the European Research Council (ERC) under the European Union’s Horizon 2020 research and innovation program (grant agreement No. 755745) and the Israel Science Foundation (380/17).

We used the significant change in activity between baseline and event, rather than significant tuning as a criterion for neuron selection to include all task-related neurons in the analysis while allowing for the accumulation of small directional effects. Selecting neurons based on significant tuning did not alter the results.

We restricted each analysis to the relevant subset of neurons. We included all the neurons in the effect size analysis (Table 1, row index A-D, overall column). Only neurons with a significant main effect in at least one of the two conditions were included in the tuning consistency score, PCA and alignment index analyses (Table 1, row index E-J, significant neurons column). Only neurons with a significant main effect in both two conditions were included in the analysis sorting PSTH by neuronal PD (Table 1, row index K-P, significant neurons column). Only neurons with a significant main effect in the two conditions were included in the population vector analysis (Table 1, row index N, significant neurons column). Including all neurons in all the analysis did not alter any of our conclusions but often resulted in more noisy estimations.

For the analysis of bin-by-bin correlations across tasks we only included neurons with significant directional tuning in both conditions using the Kruskal-Wallis test with the critical value set at 0.05. We used a lower criterion for including neurons (0.01 in the analysis described above) since the short bins reduced the power of the test. We confirmed that using the critical value of 0.01 did not alter our conclusions.

